# Strain specific persistence in the murine lung of *Aspergillus fumigatus* conidia causes an Allergic Broncho-Pulmonary Aspergillosis-like disease phenotype

**DOI:** 10.1101/2020.12.04.412726

**Authors:** Jane T. Jones, Ko-Wei Liu, Xi Wang, Caitlin H. Kowalski, Brandon S. Ross, Kathleen A. M. Mills, Joshua D. Kerkaert, Tobias M. Hohl, Lotus A. Lofgren, Jason E. Stajich, Joshua J. Obar, Robert A. Cramer

## Abstract

*Aspergillus fumigatus* is a filamentous fungus which can cause multiple diseases in humans. Allergic Broncho-pulmonary Aspergillosis (ABPA) is a disease diagnosed primarily in Cystic Fibrosis patients caused by a severe allergic response often to long-term *A. fumigatus* colonization in the lungs. Mice develop an allergic response to repeated inhalation of *A. fumigatus* spores; however, no strains have been identified that can survive long-term in the mouse lung and cause ABPA-like disease. We characterized *A. fumigatus* strain W72310 by whole genome sequencing and *in vitro* and *in vivo* viability assays in comparison to a common reference strain, CEA10. W72310 was resistant to leukocyte-mediated killing and persisted in the mouse lung longer than CEA10, a phenotype that correlated with greater resistance to oxidative stressors, hydrogen peroxide and menadione, *in vitro*. In animals both sensitized and challenged with W72310, conidia, but not hyphae, were viable in the lungs for up to 21 days in association with eosinophilic airway inflammation, airway leakage, serum IgE, and mucus production. W72310-sensitized mice that were recall-challenged with conidia had increased inflammation, Th1 and Th2 cytokines, and airway leakage compared to controls. Collectively, our studies demonstrate that a unique strain of *A. fumigatus* resistant to leukocyte killing can persist in the mouse lung in conidial form and elicit features of ABPA-like disease.

**IMPORTANCE:** Allergic Broncho-pulmonary Aspergillosis (ABPA) patients often present with long-term colonization of *Aspergillus fumigatus*. Current understanding of ABPA pathogenesis has been complicated by a lack of long-term *in vivo* fungal persistence models. We have identified a clinical isolate of *A. fumigatus*, W72310, which persists in the murine lung and causes an ABPA-like disease phenotype. Surprisingly, while viable, W72310 showed little to no growth beyond the conidial stage in the lung. This indicates that it is possible that *A. fumigatus* can cause allergic disease in the lung without any significant hyphal growth. The identification of this strain of *A. fumigatus* can not only be used to better understand disease pathogenesis of ABPA and potential anti-fungal treatments, but also to identify features of fungal strains that drive long-term fungal persistence in the lung. Consequently, these observations are a step toward helping resolve the long-standing question when to utilize antifungal therapies in patients with ABPA and fungal allergic type diseases.

## INTRODUCTION

Individuals with atopic asthma, COPD, and particularly Cystic Fibrosis (CF) are susceptible to chronic fungal colonization and infections in the lung (1). For example, 30-80% of CF patients persistently test positive for growth of the filamentous fungus, *Aspergillus fumigatus,* in expectorated sputum (2–4) and this finding is associated with overall decreased lung function (5). These findings demonstrate that *A. fumigatus* colonization is a critical aspect of CF disease pathogenesis. A subset of these patients develops Allergic Broncho-pulmonary Aspergillosis (ABPA), a particularly difficult disease to diagnose and treat. Individuals with ABPA exhibit a type 2 immune response that leads to the production of high levels of IL-4, airway eosinophilia, and increased total and *A. fumigatus* specific IgE (6, 7). Recurrent ABPA exacerbations lead to the development of bronchiectasis, airway remodeling, and fibrosis as a long-term consequence of fungal colonization (8). Although treating ABPA patients with anti-fungal drugs can improve symptoms in some cases (9, 10) it is not known how or whether persistence of *A. fumigatus* in the lungs specifically contributes to ABPA disease pathogenesis.

Murine models of fungal persistence using agar beads conjugated to *A. fumigatus* conidia successfully recapitulated long term (21 to 28 days) colonization of fungal hyphae in the lung (11, 12), however, mice in this specific agar bead model do not develop the strong IgE/Th2 response observed in ABPA patients. Numerous other mouse models that seek to recapitulate human ABPA have been described, including utilizing *A. fumigatus* antigens (13) and repeated challenge with live conidia (14), though none of these models exhibited long-term persistence of *A. fumigatus* in the airways, a common and important feature of ABPA 5, 15, 16). Importantly, the extent of fungal growth in ABPA patients’ airways is unclear, though hyphae have been identified in sputum and tissue in some patient samples (17, 18). Consequently, *in vivo* studies examining the fungal contribution to ABPA development, persistence, and disease progression are currently difficult to conduct. This is a critical question to address because it is expected to help inform when antifungal therapies may be effective in the context of ABPA.

The goal of the current study was to identify a strain of *A. fumigatus* that is capable of long-term persistence in the lung of immune competent mice. Using a clinical strain isolated from a sputum sample, viable *A. fumigatus* was recovered from the murine airways for up to 21 days. Animals with persistent fungal burden developed increased serum IgE, eosinophilia, airway damage, mucus production, and an increased immune response to re-exposure to fungi, all common features of ABPA. Surprisingly, these symptoms developed despite little to no hyphal growth observed in the airways of animals with persistent fungal burden. Consequently, these data demonstrate that a fungal strain with resistance to leukocyte killing and relatively low virulence is capable of long-term persistence in the murine lung and initiating ABPA-like disease pathogenesis.

## MATERIALS AND METHODS

### Animal Inoculations

Mice were housed in autoclaved cages at ≤4 mice per cage with HEPA-filtered air and water. For single-exposure studies, outbred Wild Type C57Bl/6 female mice 8-12 weeks old (Jackson Laboratories) were intranasally inoculated with 1×10^7 live conidia (in 40μl PBS) per mouse. *A. fumigatus* strains CEA10 and W72310 were grown on 1% glucose minimal media (GMM) for 3 days at 37°C, conidia were collected in 0.01% Tween and washed 3 times in sterile PBS. Animals were inoculated with 3×10^7 live conidia for 36 hours (FLARE), 1×10^7 live conidia 1X in week one and 5X in week two (sensitization), and 1×10^7 live conidia on the first day of week three (challenge). Animals were monitored daily for disease symptoms and we carried out our animal studies in strict accordance with the recommendations in the *Guide for the Care and Use of Laboratory Animals*. The animal experimental protocol 00002241 was approved by the Institutional Animal Care and Use Committee (IACUC) at Dartmouth College.

### Hygromycin-resistant and mRFP strain generation

Protoplasts from strains CEA10 and W72310 were generated with *Trichoderma harzianum* (Millipore Sigma) lysing enzyme and transformed ectopically with linear constructs of *gpdA*-driven *hph* hygromycin-resistance gene or H2A:mRFP (CEA10) as previously described (19, 20).

### RNA preparation and NanoString Analysis

Animals were “sensitized” to either PBS, CEA10, or W72310 as described previously and lungs were removed at euthanasia for mRNA analysis. Lungs were freeze dried, homogenized with glass beads using Mini Bead Beater (BioSpec Products Inc, Bartlesville, OK) and resuspended in Trizol (Thomas Scientific) and chloroform to extract RNA according to manufacturer’s instructions. After RNA was assessed for quality, 500ng of RNA was used per NanoString reaction using the nCounter Mouse Immunology v1 Gene Expression Panel (NanoString). nSolver 4.0 software was used for background subtraction and normalization. nCounter Advanced Analysis 2.0 was used for quality control analysis and pathway analysis. For additional pathway analysis, accession numbers were converted to Entrez IDs via the DAVID Gene Accession Conversion Tool (21, 22). The Entrez IDs were then used to pull GO terms for each gene from the Mouse Genome Informatics’ website (http://www.informatics.jax.org/batch). The resulting file was reformatted to fit the TopGO package’s requirements for gene ID to GO ID conversion (`gene_ID \t GOID1;GOID2;…;GOIDX`). Genes were classified as increased or decreased based on a 2-fold change cutoff and a p-value ≤ 0.05. Lists of all differentially expressed genes (DEGs), increased DEGs, or decreased DEGs were inserted into separate TopGOdata objects using the gene ID to GO ID conversion file to assign all possible GO terms for each gene. A nodesize cutoff of 10 was used and a Classic Fischer’s test followed by a Benjamini-Hochberg correction for multiple testing was performed via the TopGO R package (23) to determine enriched GO terms within the datasets. The degree of enrichment was calculated as the number of significant genes divided by the number of genes expected by random chance. Data was plotted via the ggplot2 and ComplexHeatmap (24) packages in R version 3.6.2 (2019-12-12).

### Colony Forming Units

Whole lungs were homogenized in 1ml sterile PBS using glass beads in a Mini Bead beater. Serial dilutions (1:10-1:1000) were then spread onto agar plates containing Sabouraud media or Sabouraud media containing 175μg/ml hygromycin and incubated O/N at 37°C. Plate dilutions containing 50-100 visible colonies were quantified and expressed as a measurement of Colony Forming Units per ml (CFUs/ml).

### Histology: GMS, PAS, H&E

After euthanasia, cannulas were inserted into trachea and lungs were excised from the body cavity. Lungs were inflated with 10% buffered formalin phosphate for 24h and stored in 70% ethanol until embedding. Paraffin-embedded sections were stained with hematoxylin and eosin (H&E), Grocott-Gomori methenamine silver (GMS), and Periodic Acid Schiff (PAS). Slides were analyzed microscopically with a Zeiss Axioplan 2 imaging microscope (Carl Zeiss Microimaging, Inc., Thornwood, NY) fitted with a QImaging Retiga-SRV Fast 1394 RGB camera.

### Serum Analysis

Cardiac punctures were performed post-mortem. After 1 hr at room temperature, blood samples were centrifuged at 2000g for 30 minutes and serum was isolated. Total IgE was measured by ELISA using kit (Invitrogen) and performed according to manufacturer’s instructions.

### Broncho-Alveolar Lavage (BAL) Analysis (inflammatory cells, Albumin, ELISAs)

After euthanasia, cannulas were inserted into trachea and lungs were removed. Lungs were lavaged with 3 sequential cold PBS washes (1ml, 0.5ml, 0.5ml). BAL was centrifuged at 300g for 5 minutes and supernatant was removed for cytokine (IL-4, IL-10, and IFNγ R&D) and albumin (Eagle Diagnostics) analysis according to manufacturer’s instructions. Remaining cells were counted and centrifuged onto slides using Rotofix Cytospin. Up to 300 cells were counted to determine percentages of macrophages, neutrophils, eosinophils, and lymphocytes. Total numbers of individual cell populations were calculated using individual percentages multiplied by total BAL cell numbers.

### Cell Death Assay

L-GMM was inoculated with 2.0 × 10^6 mRFP-CEA10 or mRFP-W72310 and incubated with either 25μM menadione or 5mM hydrogen peroxide for 6 hr at 37°C. After incubation, conidia were run on Beckman Coulter Cytoflex S Flow Cytometer for % positive RFP (viability). Analysis was performed using FlowJo version 9.9.6 as previously described (25).

### Isolation of Bone Marrow and Co-Culture Conditions

After euthanasia, femur and tibia were extracted from 8-12 week old C57Bl/6 mice and the bones were flushed with PBS to collect bone marrow-derived cells. After red blood cell lysis, cells were counted and cultured with either mRFP-CEA10 or mRFP-W72310 AF633-stained conidia as described (Jhingran et al. 2012) for 16h 10:1 MOI with 10 ng/ml interferon gamma (IFNγ) and 25mM hydroxyurea (HU).

### Flow Cytometry and Fluorescence Aspergillus REporter (FLARE) Analysis

Whole lung single cell suspensions were prepared as described (20). Briefly, lungs were minced and digested in buffer containing 2.2 mg/ml collagenase type IV (Worthington), 100μg/ml DNase I (Zymo Research) and 5% FBS at 37°C rotating for 45 minutes. Digested samples were then passed through an 18 gauge needle, resuspended in Red Blood Cell lysis buffer, diluted with PBS, passed through a 100μm filter, and counted. For flow cytometry analysis, cells were stained with a viability dye (eFluor 780, eBioScience), anti-CD45 (Pacific Orange, Invitrogen), anti-CD11b (PECy5, BioLegend for cellularity, PerCPCy5.5, BioLegend for FLARE), anti-CD11c (PE, BioLegend), anti-Ly6G (FITC, BioLegend), anti-CD64 (BV421, BioLegend) and anti-SiglecF (BV421, BD bioscience). Samples were analyzed on a MacsQuant VYB flow-cytometer (cellularity) or Beckman Coulter Cytoflex S (FLARE). Macrophages were identified as CD45^+^/Ly6G^−^/CD11b^+^/CD64^+^, neutrophils were identified as CD45^+^/SiglecF^−^/Ly6G^+^/CD11b^+^ and eosinophils were identified as CD45^+^/SiglecF^+^/CD11c^low^. Analysis was performed with FlowJo version 9.9.6.

### Genome Sequencing and Variant Analyses

Mycelial cultures of *A. fumigatus* using liquid glucose minimal media with yeast extract were inoculated in small petri dishes grown overnight (18-24hrs) at 37C. Mycelia were collected, lyophilized, and bead beaten to powder and DNA was extracted as previously described (27). Genomic sequencing libraries of the DNA were constructed using the SeqOnce (SeqOnce Biosciences, Pasadena, CA) with multiplexing barcodes by the Genomics Core at UC Riverside Institute for Integrative Genome Biology. The genomic libraries were sequenced as 2 x 150 bp reads on Illumina Novaseq 6000 (Illumina, San Diego, CA) at UC San Francisco Sequencing Core to generate ~2.17 Gb of sequence. Sequence data for W72310 was deposited in the Sequence Read Archive under BioProject PRJNA614926. Sequence data for strain CEA10 was obtained from Sequence Read Archive under accession ERR232423 and BioProject PRJEB1497. The sequence reads for each strain were aligned to the Af293 genome downloaded from FungiDB v.46 (28, 29) using BWA v0.7.17 (30) and converted to the BAM file format after running fixmate and sort commands in samtools v1.10 (Li et al., 2009). Duplicate reads were removed and reads indexed using MarkDuplicates and Build BamIndex in picard tools v2.18.3 (http://broadinstitute.github.io/picard). To avoid overcalling variants near alignment gaps, reads were further realigned using RealignerTargetCreator and IndelRealigner in the Genome Analysis Toolkit GATK v3.7 (Auwera et al., 2013). The variants (SNPs and INDELS) for W72310 and CEA10 were genotyped relative to the *A. fumigatus* reference genome Af293 using HaplotypeCaller step in GATK v4.0 (doi: 10.1101/201178). Filtering was accomplished using GATK’s SelectVariants call with the following parameters: for SNPS: -window-size = 10, -QualByDept ≤ 2.0, -MapQual ≤ 40.0, - QScore ≤ 100, -MapQualityRankSum ≤ -12.5, -StrandOddsRatio > 3.0, - FisherStrandBias > 60.0, -ReadPosRankSum ≤ −8.0. For INDELS: -window-size = 10, - QualByDepth ≤ 2.0, -MapQualityRankSum ≤ −12.5, -StrandOddsRatio > 4.0, - FisherStrandBias > 200.0, -ReadPosRank ≤ -20.0, -InbreedingCoeff ≤ −0.8. Resultant variants were annotated with snpEff (Cingolani et al., 2012) using the Gene File Format gene annotation for Af293 v.46 in FungiDB. Variants that overlapped transposable elements (TEs) were removed by positional mapping to locations of annotated TEs in the FungiDB v.46 release of the Af293 reference genome, using BEDtools -subtract (Quinlan and Hall, 2010). Mutations in W72310 were analyzed relative to CEA10 using a custom script implemented in the R programing environment (R core team 2013). To identify mutations occurring in allergen genes, a database of genes putatively capable of eliciting immune response was curated from the published literature (n = 113) (Table 2) and mapped against W72310 variant positions using a custom script in R. Putative allergen genes in W72310 containing variants other than synonymous or intergenic mutations were annotated in Fungi DB v.46 (29). To investigate variants unique to W72310 and occurring in genes relevant to the phenotypes of interest in this study (oxidative stress, cell wall integrity, primary metabolism, and melanin production), we used reverse GO mapping of the terms GO: 0006979 (response to oxidative stress), GO:0005618 (cell wall), GO:0044238 (primary metabolic process) GO:0006582 (melanin metabolic process) and GO:0042438 (melanin biosynthetic process). The code and datafiles for variant assessment associated with this project can be accessed via GitHub in the repository: stajichlab/W72310 (DOI: 10.5281/zenodo.4116457)(31).

### Statistical Analysis

All statistical analyses were performed with Prism 8.3 software (GraphPad Software Inc., San Diego, CA). For *in vitro* comparison of 2 groups, Student’s T-test was used. For animal experiments, nonparametric analyses were performed (Kruskal-Wallis, Dunn’s multiple comparisons; Mann-Whitney, single comparisons). All error bars represent standard errors of the means. NS P>0.05; * P≤0.05; ** P≤0.01; *** P≤0.001; **** P≤0.0001.

## RESULTS

### Aspergillus fumigatus clinical isolate, W72310, is resistant to immune cell-mediated killing and persists in the murine lung

Previous work from our laboratory found that a clinical isolate strain, W72310, had reduced virulence compared to the commonly studied reference strain, CEA10, in mice with suppressed immune systems (32). Conidia from W72310 have also been reported to germinate more slowly under a variety of conditions (33) in comparison to other *A. fumigatus* strains. We hypothesized that a slow growing, less virulent strain may evade immune clearance and allow establishment of long-term fungal persistence in murine airways. In order to test this hypothesis, we first compared W72310 and CEA10 persistence in the immune competent murine lung 7 days after one fungal intranasal inoculation. GMS staining of fixed lungs revealed that fungal conidia were still present in W72310-exposed lungs 7 days post inoculation, however, no observable conidia were found in CEA10 challenged animals (Fig. 1A). While swollen conidia could be observed in W72310 inoculated animals, no hyphal growth was visible across multiple independent histopathology sections. In a parallel experiment, quantification of total colony forming units revealed viable fungi only from lungs inoculated with W72310 (Fig 1B), indicating W72310 persists longer than CEA10 in C57BL6/J mice.

**Figure 1:**
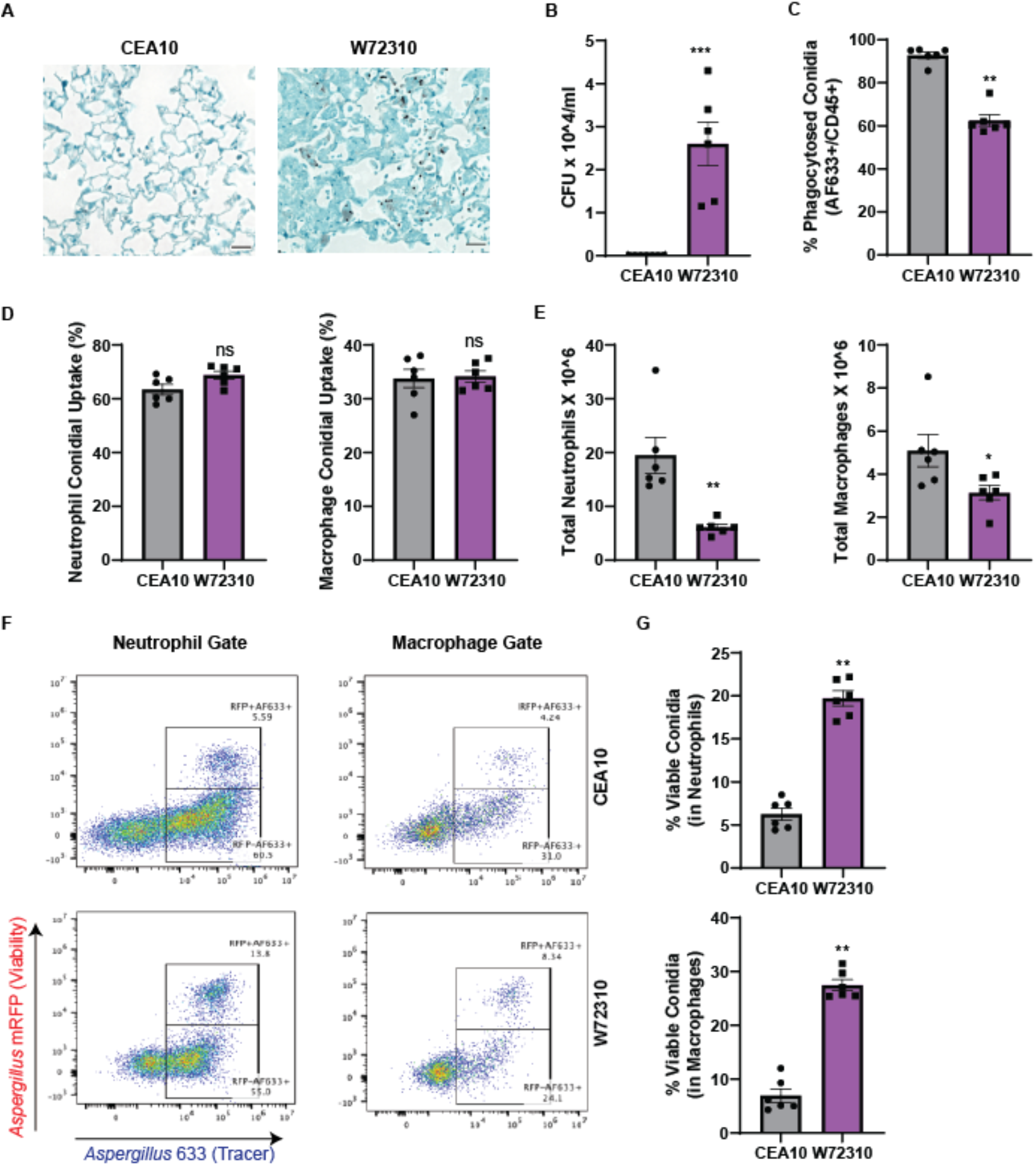
*A. fumigatus* clinical isolate, W72310, persists longer in the murine lung and is resistant to leukocyte-mediated death compared to common laboratory strain, CEA10. **(A)** C57Bl/6 mice were intranasally inoculated with 1X10^7 live CEA10 or W72310 conidia and euthanized after 7 days. Lungs were fixed in formalin and stained for fungi (GMS stain, black). N=6-8 animals, scale bars=100um, 40X. **(B)** Lungs from parallel experiments were assessed for total colony forming units (N=6-8 animals). C57Bl/6 mice were oropharyngeally inoculated with 3.0X10^7 AF633-stained mRFP-CEA10 or mRFP-W72310 conidia for 36hrs. Lungs were analyzed by flow cytometry for **(C)** % phagocytosis of conidia in immune cells (AF633^+^/CD45^+^), **(D)** % positive conidia (AF633) in neutrophils and macrophages, and **(E)** total neutrophils (CD45^+^/Ly6G^+^/CD11b^+^) and macrophages (CD45^+^/Ly6G^−^/CD11b^+^/CD64^+^). **(F)** Viability of conidia phagocytosed in neutrophils and macrophages was analyzed by flow cytometry and **(G)** quantified. Representative micrographs were selected out of 6 mice per group. Mann-Whitney single comparisons were used and NS P>0.05; * P≤0.05; ** P≤0.01; *** P≤0.001.

To address how W72310 conidia persist in the lung, we assayed the susceptibility of W72310 and CEA10 fungal conidia to killing by bone marrow-derived neutrophils and macrophages *ex vivo*. CEA10 and W72310 that ectopically express mRFP were generated in order to distinguish live (mRFP^+^) from dead (mRFP^−^) conidia by flow cytometry. Given the relatively short half live of RFP (approximately 45 minutes) in the phagolysosome of leukocytes, this fluorescent protein functioned as a fungal viability marker during cellular interactions with leukocytes. A fluorescent tracer dye (Alexa Fluor 633) was covalently attached to the surface of the mRFP-expressing strains and served as a conidial tracer (26). After co-culture of either mRFP-W72310/AF633 or mRFP-CEA10/AF633 conidia and bone marrow cells for 16 h, there were 15% more neutrophils positive for CEA10 conidia compared to W72310 conidia, while there were no differences observed in macrophages (Supp fig. S1A). A small but significant increase in survival of W72310 was observed in neutrophils compared to CEA10. The survival of W72310 conidia was higher by 15% compared to CEA10 in macrophages (Supp fig. S1B). To test this observation *in vivo*, we utilized the FLARE assay to quantify leukocyte uptake and viability of W72310 and CEA10 conidia in the murine lung. 36 hr after inoculation with AF633-stained mRFP-CEA10 or W72310 conidia, the percentage of W72310 conidia that were phagocytosed by CD45^+^ immune cells was 30% less than the percentage of CEA10 conidia in CD45^+^ immune cells (Fig. 1C). However, comparison of neutrophil and macrophage conidial uptake (% positive AF633) showed no difference between CEA10 and W72310 (Fig. 1D), in contrast to observations in the bone marrow-derived neutrophils. The reduced phagocytosis of W72310 conidia correlated with a four-fold reduction in neutrophils and two-fold reduction in total macrophages in the lungs of mice inoculated with W72310 compared to CEA10 (Fig. 1E). The percentage of viable W72310 conidia phagocytosed by neutrophils and macrophages significantly increased four-fold in comparison to CEA10 (Fig. 1F and G). Taken together, these data suggest the *A. fumigatus* strain W72310 recruits less inflammatory cells and is killed less efficiently than the highly virulent CEA10 strain in a murine model of fungal bronchopneumonia.

### Comparison of W72310 sensitization to CEA10 sensitization in the murine lung

Given the persistent presence of the W72310 strain in the murine lung, its increased resistance to leukocyte-mediated killing, and the altered cellular response to W72310 conidia, we next compared the overall immune response to sensitization with CEA10 and W72310 live conidia. Animals were intranasally inoculated and sensitized to W72310 or CEA10 conidia as indicated (Fig. 2A). Compared to PBS-sensitized animals, CEA10 and W72310 animals had 6- and 8-fold increased serum IgE, respectively (Fig. 2B). Although we observed a trend for more total IgE in animals exposed to W72310 compared to CEA10, the difference was not statistically significant. Additionally, small but detectable CFUs were observed in lungs sensitized with CEA10, however, approximately 10^6 CFUs were detected in lungs of animals sensitized with W72310 (Fig. 2C). In parallel, total RNA was extracted from whole lungs and analyzed for changes in mRNA levels of immune-related genes using NanoString nCounter technology. As expected, overall changes in mRNA levels were strikingly different in PBS-sensitized animals compared to the two fungal strains (Fig. 2D). Increased signatures in pathways related to cytokine/chemokine signaling, host pathogen interactions, and innate and adaptive immune signaling were observed in fungal challenged animals as expected (Fig. 2E.). Interestingly, the only immune-related pathways with reduced mRNA levels in fungal-exposed mice compared to PBS were TGF-β and inflammasome encoding genes (Fig. 2E). Using GO-term analysis with a cut-off of P≤0.01 to differentiate highly significant pathways in the W72310-sensitized lung compared to PBS, analysis showed prevalent eosinophil, monocyte, and lymphocyte chemotaxis, ERK1 and ERK2 signaling, IFNγ-response signaling, and general immune system responses (Fig. 2F). Comparison of PBS and W72310- sensitized lungs revealed 174 increased and 38 decreased mRNA transcripts (Fig. 2G). Surprisingly, transcripts of only 9 genes were differentially abundant (2-fold or higher) between CEA10 and W72310 challenged mice. Of these, transcripts of all 9 genes were increased in W72310 sensitized animals compared to CEA10 sensitized animals (Fig. 2H). These differentially expressed transcripts are encoded by the genes *nos2, clec4e, cxcl-9, cxcl-10, ccl4, cxcl-11, klra21, cxcl3, and tnfα*; transcripts largely involved in T cell/monocyte recruitment and activation, host defense, and CXCR3 receptor activation. The lack of differences between CEA10 and W72310 challenged animals seems especially striking given the clear differences in persistent fungal burden.

**Figure 2:**
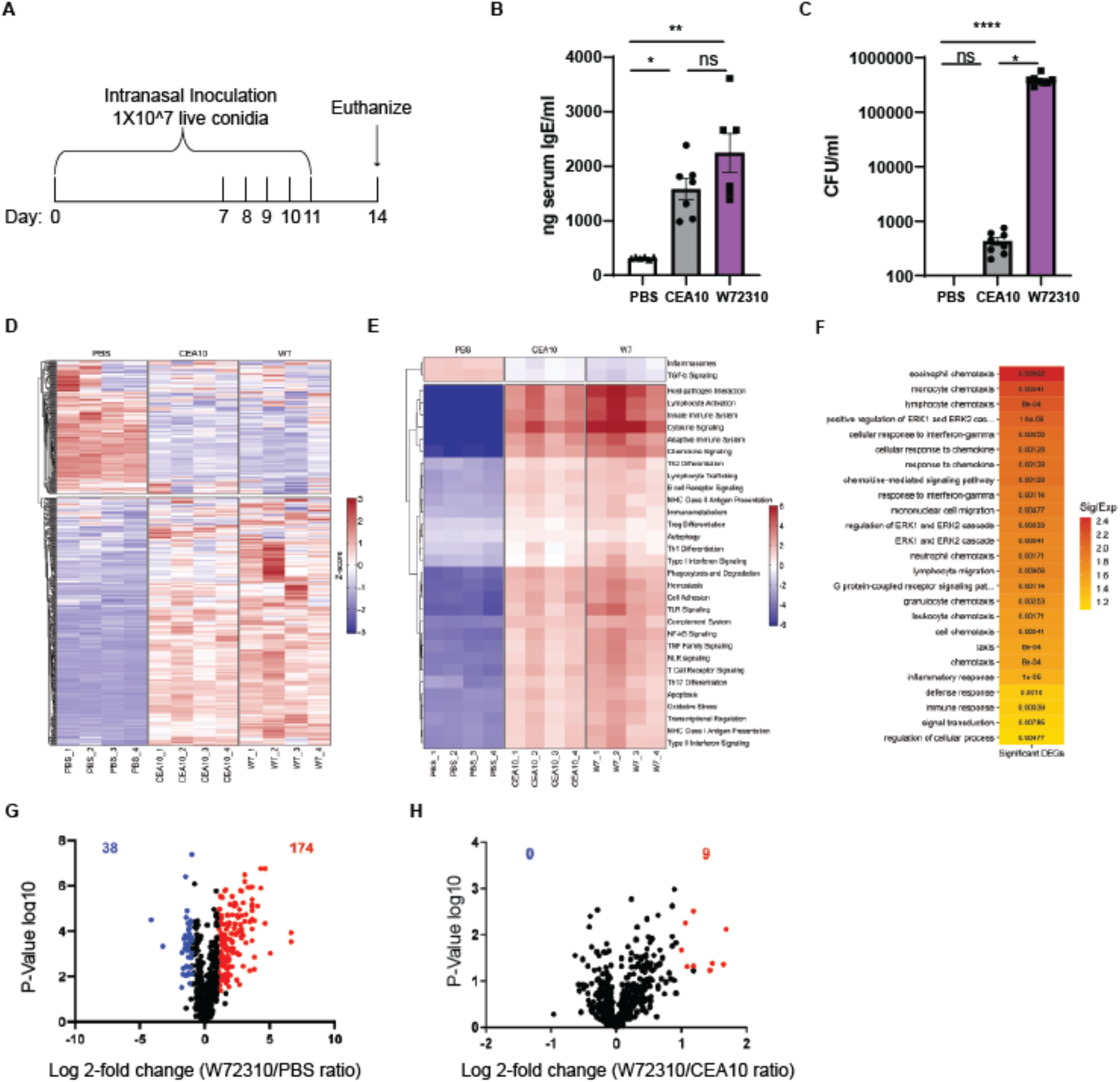
W72310-induced sensitization is similar to CEA10-induced sensitization in the murine lung. **(A)** Mice were inoculated with either PBS or 1X10^7 live W72310/CEA10 conidia on the indicated days and euthanized on day 14. **(B)** Total serum IgE was measured by ELISA on blood samples collected from animals. N=6-7 mice. **(C)** Total CFUs were measured in the lungs of mice. N=8 mice. Mann-Whitney with Dunn’s multiple comparison were used and * P≤0.05; ** P≤0.01; *** P≤0.001; **** P≤0.0001. **(D)** Total mRNA was extracted from the whole lung and gene expression was analyzed by NanoString nCounter^®^ Immunology panel. Downregulation (blue) or upregulation (red) of individual replicates were represented by heat map, n=4 mice/group. **(E)** Pathway signature of mice sensitized to either PBS, CEA10 or W72310. **(F)** Heatmap showing the degree of enrichment and p-value for each GO-term enriched in increased genes. The color of each tile indicates the log2 transformed degree of enrichment for a given term within each gene set. The p-value is overlaid in text to show the significance of each term. The terms on the y-axis are ordered according to decreasing degree of enrichment from top to bottom. **(G)** Volcano plot showing the distribution of fold changes in gene expression in W72310-treated mice compared to PBS: genes with absolute fold changes ≥2 and with p value ≤ 0.05 (Student’s T-test) are shown in blue and genes with absolute fold changes ≥2 and with p value ≤ 0.05 (Student’s T-test) are shown in red. **(H)** Volcano plot showing the distribution of fold changes in gene expression in W72310-treated mice compared to CEA10: genes with absolute fold changes >2 and with p value ≤ 0.05 (Student’s T-test) are shown in red.

### W72310 accumulates in the murine lung during sensitization

ABPA patients are sensitized to *A. fumigatus* antigens (34). A previous report demonstrated that fungi could persist longer in an allergen-sensitized murine lung compared to a control lung (35), so we next determined whether sensitization with W72310 or CEA10 increased persistence of a subsequent exposure to W72310 or CEA10 conidia. In light of the data from the immune competent murine model (Fig. 1), it is likely that W72310 sensitization alone would cause an accumulation of fungi in the lung, so in order to differentiate between fungi present from sensitization and fungi present due to challenge, we generated CEA10 and W72310 strains that were resistant to hygromycin. Mice were sensitized with either PBS, WT CEA10 or WT W72310 and challenged with either PBS, W72310-hyg or CEA10-hyg. Colony forming units were measured on Sabouraud Dextrose Agar (SAB)-containing plates (total CFUs) or SAB/hygromycin-containing plates (“challenge” CFUs) from the lungs of mice euthanized 7 days after challenge (Fig. 3A). Interestingly, no live conidia were detected in mice sensitized with CEA10 and challenged with PBS, demonstrating that repeated intranasal (IN) inoculation was not sufficient to drive CEA10 fungal conidia accumulation in the murine lung (Fig. 3B). In contrast, approximately 1X10^7 live conidia were detected from mice sensitized with W72310 and challenged with PBS, indicating an accumulation of viable fungi in the lung from repeated W72310 IN sensitizations (Fig. 3B). Challenge with CEA10-hyg in W72310-senstized mice did not facilitate persistence of CEA10, however, an almost two-fold increase in CFUs was observed in W72310-challenged mice when sensitized with W72310 but not CEA10 (Fig. 3B). Collectively these data indicate that sensitization causes a modest increase in persistence of W72310 but not CEA10. Consequently, fungal sensitization itself is not sufficient nor necessary to promote fungal persistence in the lung with all strains of *A. fumigatus*.

**Figure 3:**
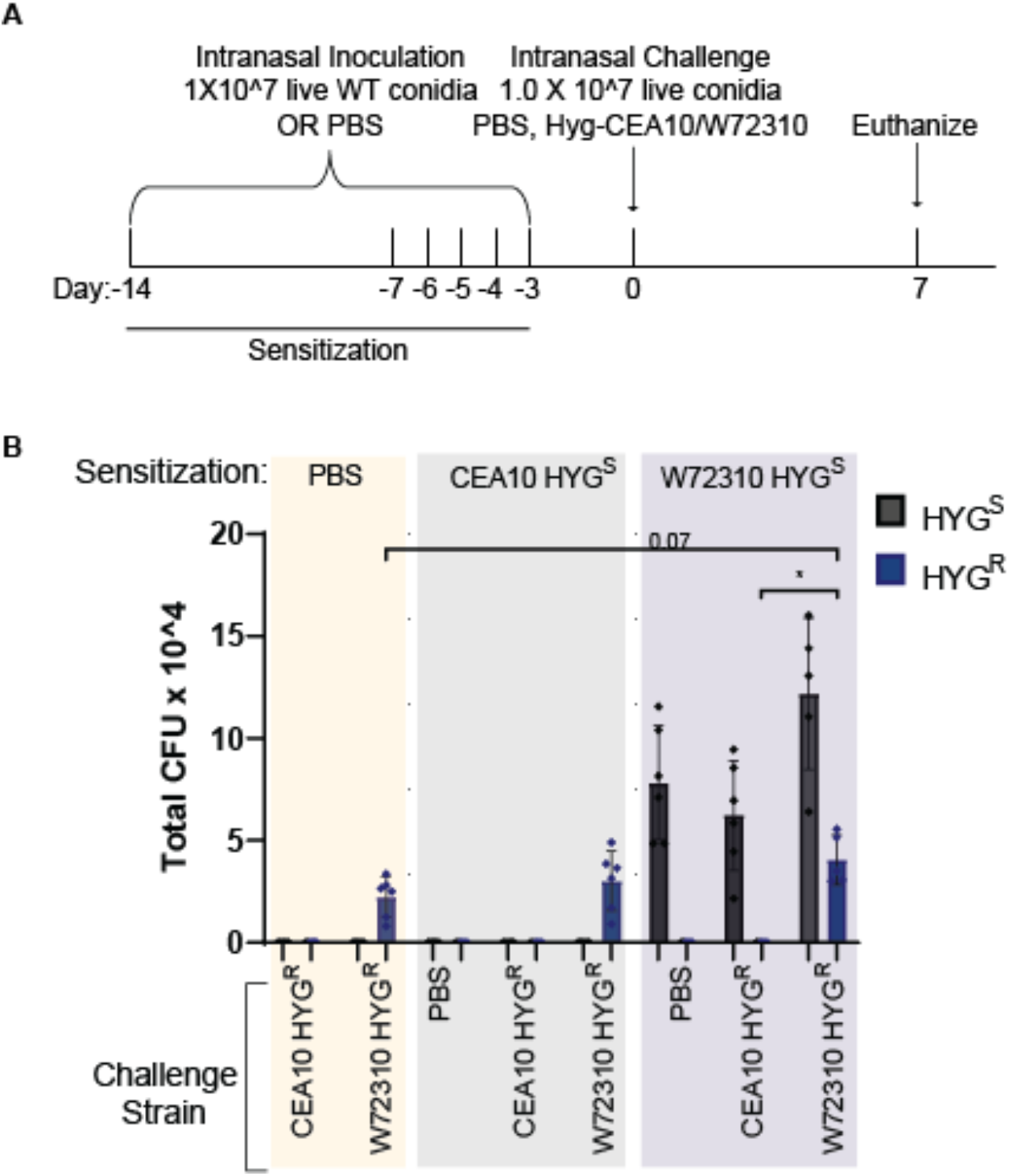
W72310 accumulates in the mouse lung during sensitization. **(A)** C57Bl/6 mice were inoculated with either PBS or 1X10^7 live W72310/CEA10 conidia on the indicated days (sensitization) and challenged on day 0 with either PBS, hyg-CEA10, or hyg-W72310; animals were euthanized 7 days after challenge. **(B)** Lung homogenates were spread on plates containing SAB alone or SAB plus hygromycin and quantified for total CFU’s. “Challenge” CFUs were identified as colonies grown on hygromycin plates and “Sensitization” CFUs were calculated by subtracting hygromycin CFU values from total CFU values (SAB alone plates). HYG^S^=hygromycin sensitive and HYG^R^= hygromycin resistant. N= 6-12 mice per group. Mann-Whitney with Dunn’s multiple comparison were used and * P≤0.05.

### Sensitization and Challenge with W72310 causes increased fungal persistence in the lung in association with increased serum IgE

To determine whether W72310 persists long-term in the mouse lung, we sensitized animals as described in Figure 2 and subsequently “challenged” mice with another fungal dose on day 0 (Fig. 4A). As a control, animals were sensitized to PBS only and challenged with one dose of W72310 on day 0. To determine fungal persistence, CFUs were chosen for quantitation despite the filamentous morphology of *A. fumigatus* because hyphal elements could not be observed in histopathology samples at any time point. Total CFUs increased in the lungs of animals sensitized and challenged with W72310 at 2, 7, and 21 days post challenge in comparison to PBS-sensitized controls (Fig. 4B). This significance was not observed in lungs 28 days post challenge. CFUs were detected 2 and 7 days post challenge in the PBS sensitized control group. Surprisingly, CFUs were still detectable 21 and 28 days post challenge in this group as well, although to a much lesser extent. Fixed lungs were stained for fungi (GMS) to observe the fungal presence and morphology. At 2 and 7 days post challenge in both PBS and W72310-sensitized mice, prominent conidia were visible (Fig. 4C). Despite the quantification of detectable CFUs in PBS-sensitized mice 21 and 28 days post challenge, conidia were not visible by GMS stain (Fig. 4C). Conidia were strongly visible 21 days after challenge in the W72310- sensitized mice, however, they appeared to be taken up in large vacuolar phagocytes (Fig. 4C). Interestingly, multiple conidia were visible in a single host cell. Conidia were mostly non-detectable by 28 days, with a small number visible by GMS stain. Collectively these data demonstrate that W72310 can persist in a fungal sensitized murine lung up to one month after inoculation.

**Figure 4:**
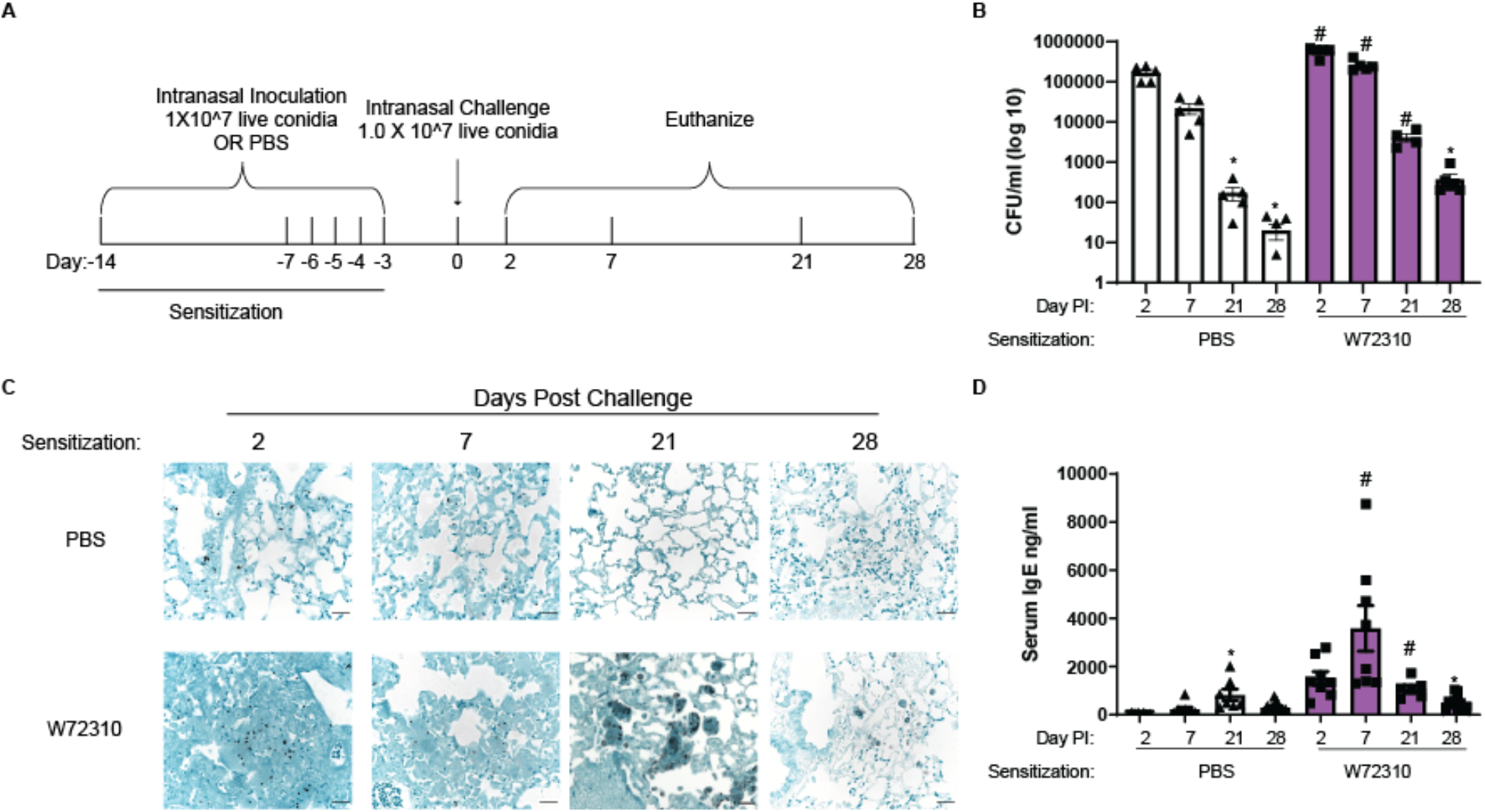
Sensitization and Challenge with W72310 causes increased fungal persistence in the lung in association with increased serum IgE. **(A)** Schematic of ABPA model protocol: C57Bl/6 mice were inoculated with either PBS or 1X10^7 live W72310 conidia on the indicated days and challenged on day 0. Animals were subsequently euthanized on days 2, 7, 21, and 28. **(B)** Colony Forming Units (CFUs) were quantified from total lung homogenates at the indicated time points and represented on a log scale. **(C)** Paraffin-embedded fixed lungs were stained for GMS and images were captured at 40X, scale bar=100μm. **(D)** Total serum IgE was measured by ELISA on blood samples collected from animals. (B and D) Kruskal Wallis with Dunn’s comparison to 2 day time point (within same sensitization protocol) *≤ 0.05; Kruskal Wallis with Dunn’s comparison to same time point (within different sensitization protocol) #≤ 0.05. Data include 2 independent experiments and n=8-12 mice.

Blood was collected from each animal and total serum IgE levels were measured by ELISA to determine the overall allergic response. As expected, total IgE levels increased 2 and 7 days post challenge in W72310 sensitized mice compared to controls (Fig. 4D), however significant increases were not detected at 21 and 28 days. Interestingly, even in the PBS-sensitized animals, total serum IgE was increased at 21 days compared to PBS-sensitized control at 2 days. Overall, these data indicate that sensitization and challenge with strain W72310 maintain high levels of IgE.

### Sensitization and Challenge with W72310 causes increased inflammation, leakage, and mucous cell metaplasia

Eosinophilia is a hallmark of ABPA disease (36). Differential staining of airway/bronchoalveolar lavage cells revealed that eosinophils were increased 2, 7, and 21 days after sensitization and challenge with W72310 live conidia in comparison to PBS-sensitized controls (Fig. 5A). Moreover, macrophages and lymphocytes were increased at all time points (Fig. 5A). Interestingly, lymphocytes and eosinophils continued to be detectable 28 days post challenge, but at lower levels. Overall neutrophils decreased over time, but no significant differences were observed between control and experimental groups at any time points (Fig. 5A). The inflammatory response was also assessed by staining lung sections for H&E. Some cellular infiltrates were observed in PBS-sensitized mice 2 and 7 days post challenge with W72310, while robust inflammation, including eosinophils, was observed in animals sensitized and challenged with W72310 at all time points (Fig. 5B). Furthermore, animals sensitized and challenged with W72310 for all time points had detectable mucous cells in airways as shown by PAS-stained lung sections (Fig. 5B, black arrows). At early time-points, 2 and 7 days after challenge, there were a large number of mucus-producing cells, which persisted at lower levels at later time-points.

**Figure 5:**
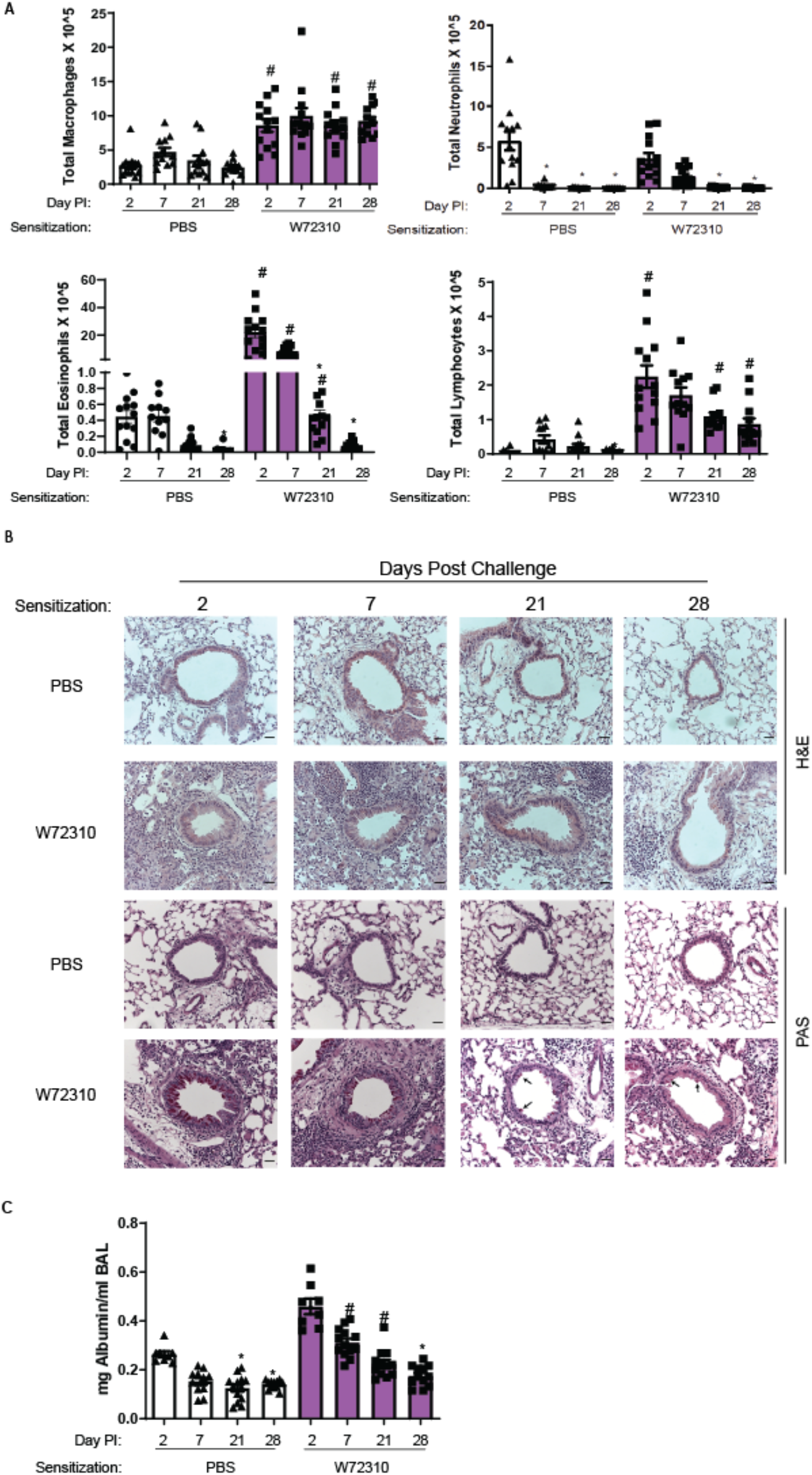
Sensitization and Challenge with W72310 causes increased pulmonary inflammation, leakage, and mucous cell metaplasia. **(A)** Total BAL was quantified for individual cell populations of macrophages, neutrophils, lymphocytes, and eosinophils. **(B)** Paraffin-embedded fixed lungs were stained for H&E and PAS and images were captured at 20X, scale bar=100μm. **(C)** BAL supernatant was analyzed for total concentrations of albumin. (A and B) Kruskal Wallis with Dunn’s comparison to 2 day time point (within same sensitization protocol) *≤ 0.05; Kruskal Wallis with Dunn’s comparison to same time point (within different sensitization protocol) ≤ 0.05. Data include 2 independent experiments and n=8-12 mice.

Total albumin in the BAL was measured to evaluate the extent of vascular leakage and potential damage to the lungs in the airways. Quantification of albumin showed significant increases in animals sensitized with W72310 and challenged 7 and 21 days compared to PBS-sensitized controls, indicating airway damage that resolved by day 28 (Fig. 5C).

### Mice recall challenged with W72310 have increased airway inflammation and damage in the lungs

Major features of ABPA disease are periods of remission and exacerbation (37). To determine if mice with persistent W72310 in the lungs were susceptible to a recall response, mice were sensitized and challenged with either PBS or W72310 and 21 days after challenge all animals were recall challenged with either PBS or W72310 live conidia (Fig. 6A). BAL and lung suspension neutrophils and eosinophils were significantly increased in the W72310 recall challenged group compared to PBS recall challenged group (Fig. 6B). Serum IgE levels were increased 5-fold in mice sensitized and challenged with W72310 at 21 days compared to PBS sensitized challenged mice, as would be expected (based on results from the time course (Figs. 4D)) and BAL albumin was measured in order to assess damage and leakage in the airways (Fig. 6C). Total albumin levels were increased two-fold in the airways of mice sensitized, challenged, and recall challenged with W72310 compared to both PBS controls. Th2 cytokines IL-4/IL-10 and Th1 cytokine IFNγ were significantly increased in BAL of mice sensitized, challenged, and recall challenged with W72310 compared to PBS controls (Fig. 6D). IL-17A cytokine levels were not detectable (data not shown). Collectively these data demonstrate that W72310 persists for several weeks in a fungal-sensitized murine lung and elicits strong inflammation, damage, mucus production, and an exacerbation-like immune response.

**Figure 6:**
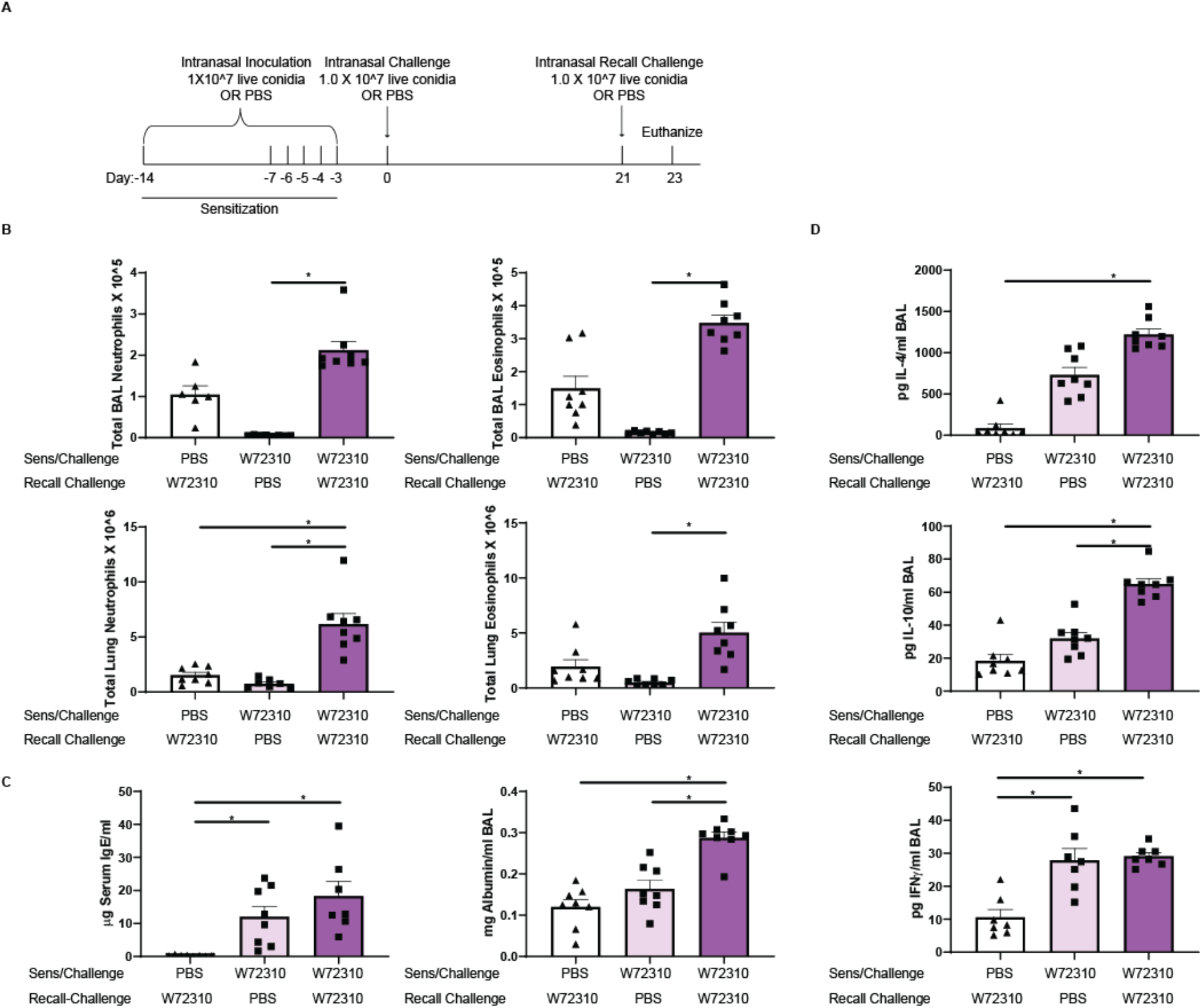
Sensitization and Challenge with W72310 and subsequent recall challenge causes increased tissue inflammation, damage, Th1/2, and IgE response. **(A)** Schematic of ABPA exacerbation protocol: C57Bl/6 mice were inoculated with either PBS or 1X10^7 live W72310 conidia on the indicated days (sensitization and challenge) and recall challenged 21 days after initial challenge. Animals were subsequently euthanized 48h later. **(B)** Neutrophils (CD45^+^/SiglecF^−^/Ly6G^+^/CD11b^+)^ and eosinophils (CD45^+^/SiglecF^+^/CD11c^low^) were quantified by flow cytometry in BAL and lung suspensions. **(C)** Serum was isolated from mice and total IgE levels were quantified and BAL supernatant was analyzed for total albumin. **(D)** BAL supernatant IL-4, IL-10 and IFNγ were measured. Mann-Whitney with Dunn’s multiple comparison were used and * P≤0.05.

### Comparison of W72310 and CEA10 whole genome sequences

Given the clear immune response differences to CEA10 and W72310, we wanted to address potential fungal mediated mechanisms for these observations. As a first step toward understanding the fungal contribution to these responses, whole genome sequencing of CEA10 and W72310 was conducted. Sequencing and variant analysis revealed a total of 56,036 variants in W72310 that differed from the AF293 reference genome, compared to only 13,014 variants identified in CEA10. These variants included 2392 positions in W72310, and 874 variants in CEA10 for which no allele could be determined (and which possibly represent a deletion at that position when compared to AF293 reference genome). We consequently identified 49,806 variants that were exclusive to W72310 and not shared with CEA10 (18,088 excluding synonymous and intergenic variants), compared to only 8302 variants found in CEA10 but not present in W72310 (1922 excluding synonymous and intergenic variants). W72310 also had a greater number of missense variants with a total of 6486, compared to only 814 in CEA10. Given the differences observed between W72310 and CEA10 in conidia viability, germination rate, and persistence in the lung, we narrowed our focus to non-synonymous variants unique to W72310 (not present in CEA10) occurring in genes involved in oxidative stress responses, cell wall biology, melanin, and metabolism using reverse GO term mapping. Variants were identified in 20 unique oxidative stress response genes, 6 cell wall genes, and 6 melanin genes (Table 1). Interestingly, no variants were identified in core metabolism genes. When non-synonymous variants unique to W72310 were mapped onto putative allergen genes, 184 SNPs and INDELS in 69 unique genes, with mutation load ranging from 1 to 21 variants per gene were found (Table 2). In order to corroborate differences observed in oxidative stress resistance, mRFP-CEA10 and mRFP-W72310 strains were assayed for susceptibility to oxidative stress inducing agents, hydrogen peroxide and menadione. The percentage of viable CEA10 conidia (RFP^+^) significantly reduced by 50% (hydrogen peroxide) and 40% (menadione), respectively. In contrast, incubation of W72310 with hydrogen peroxide and menadione only decreased the percentage of viable conidia by less than 5% (Fig. 7A). Collectively these data show that W72310 conidia are less susceptible to oxidative stress-induced death compared to CEA10 and this may contribute to its persistent phenotype in the mouse lung. The allele(s) driving this phenotype in W72310 conidia remain to be determined in future studies.

**Figure 7:**
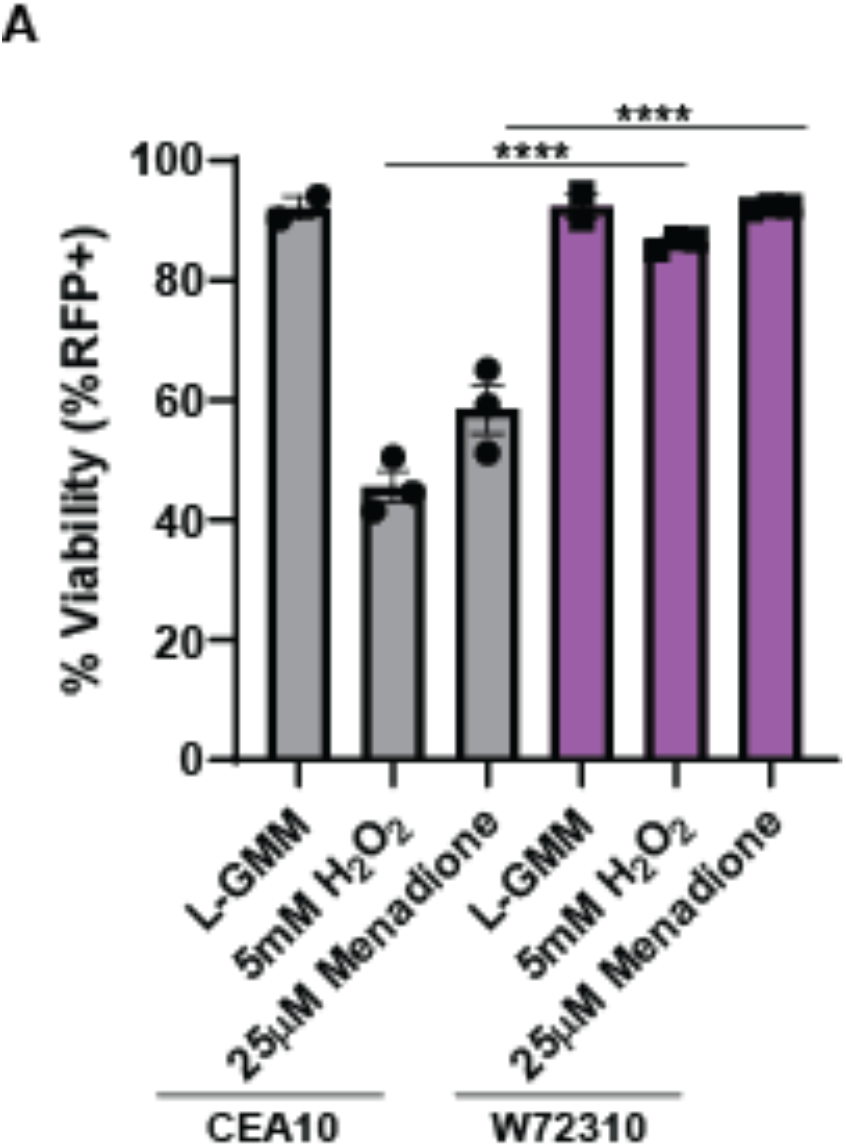
W72310 is more resistant to oxidative stress-induced death than CEA10. **(A)** mRFP-CEA10 and mRFP-W72310 were incubated with either 25μM menadione or 5mM H_2_O_2_ for 6h and analyzed by flow cytometry for viability (%RFP^+^). Data are representative of 3 independent experiments with 3-4 technical replicates/experiment. One way ANOVA with Tukey’s multiple comparison were used and **** P≤0.0001.

## DISCUSSION

In the current study, we characterized a unique clinical strain of *A. fumigatus* which persists in the murine lung and elicits an ABPA-like disease state. By identifying and characterizing the W72310 strain, our studies demonstrated that an *A. fumigatus* strain can be resistant to immune cell-mediated killing and that it can persist in the murine lung as viable conidia while eliciting a strong IgE/inflammatory/damage response. These data are important because they suggest *A. fumigatus* strains exist in the population that can persist in mammalian airways and induce pathological immune responses without robust fungal growth. Moreover, our data provide an opportunity to uncover novel fungal contributions to allergic-like diseases due to the long-term persistence of W72310 conidia in murine lungs.

We hypothesize that a key fungal mechanism for fungal persistence is enhanced oxidative stress resistance of the W72310 conidia. Reactive Oxygen Species (ROS) generated by NADPH-oxidases in neutrophils and macrophages are critical components in host defense against *A. fumigatus* infection (25, 38). Intriguingly, our data demonstrate that W72310 conidia are more resistant to both hydrogen peroxide and menadione-induced death compared to CEA10 (Fig. 7). Moreover, neutrophils and macrophages have reduced killing of W72310 conidia compared to CEA10 conidia (Fig. 1G). Genome sequencing revealed that W72310 has 20 oxidative stress response genes with high impact SNPs (Table 1). Of particular interest perhaps the catalase Afu6g03890/*catA* has 6 variants in W72310 and not CEA10 (Table 1). Previous studies demonstrated that a catalase mutant generated in *A. fumigatus* was more sensitive to hydrogen peroxide. Interestingly, its conidia did not have any difference in susceptibility to immune cell mediated killing and had similar pathogenicity in mouse models of aspergillosis compared to wild-type controls (39–41). Since there are 20 oxidative stress response genes with variants in the W72310 strain, it seems likely that more than one gene/mechanism is involved in its defense against ROS-induced killing. However, we cannot rule out that novel alleles of genes like *catA* encode proteins with increased activity.

Another fungal-centric striking observation from our studies is that W72310 persists in the mouse lung largely as viable conidia. Based on the histology at 21 days post challenge (Fig. 4C), we observed that the majority of conidia appear to be phagocytosed by large macrophage-like cells. Previous histological analyses of invasive aspergillosis patients have also demonstrated the presence of multiple conidia per giant cell (42), however, the viability of the fungus is not known. Case reports often show *A. fumigatus* in the lung as germlings or hyphae by GMS stain (43), but it is not known whether conidia, germlings, hyphae, or some combination contribute to ABPA and fungal allergic disease pathogenesis. Given that W72310 has previously been shown to have reduced germination rates in comparison to other strains (32, 33), another explanation for its persistence is that remaining in its conidial form does not allow full immune recognition of the fungus. Of note, W72310 elicited less of an inflammatory response compared to CEA10 in an immune competent pulmonary challenge model (Fig. 1E). LC3-associated phagocytosis has been shown to be critical in fungal killing and is dependent on exposure of the fungal cell wall components, melanin and β-glucan (44). Sequence analysis showed high impact SNPs in 6 cell wall genes and 6 melanin genes which could potentially prevent full recognition and/or killing of the fungus (Table 1). Collectively, SNPs in melanin and cell wall stress genes may affect overall germination and immune cell recognition of W72310 and SNPs in oxidative stress response genes could affect overall viability of phagocytosed W72310 conidia. Rigorous fungal genetic studies will be needed to determine specifically which gene(s) are important overall for the persistent phenotype.

Despite a similar immunological response to sensitization between W72310 and CEA10, select interferon-responsive genes were differentially expressed. CXCR3 was not differentially expressed between W72310 and CEA10 (data not shown), however, expression of its ligands, CXCL-9 and CXCL-10, was higher in W72310-sensitized lungs compared to CEA10. CXCL9 and CXCL10, which have been shown to be induced in other mouse models of asthma and *A. fumigatus*-induced inflammation (45–47), activate Th1-type immune responses through CXCR3. Because CXCR3 is preferentially expressed in Th1 cells, (48), this could indicate a potential role for the Th1 immune response in the overall persistence observed from W72310. In both a mouse model of severe asthma and in humans with severe asthma, CXCL-10 was shown to play a critical role in corticosteroid resistance and Th1-mediated inflammation (49). However, differences in the immune response at this time point could be either in response to the different strains of fungi or different quantities of fungi since we observe significantly more W72310 CFUs at this time point (Fig. 2C).

The results of our present study indicate that persistent conidial colonization in the fungal-sensitized mouse lung causes many features of APBA-like disease (Figs 4–6). Diagnosis of ABPA has proven to be challenging and it has long been postulated that ABPA rates are under-represented due to a lack of consistency in diagnostic methods (34). Elevated total and *A. fumigatus* specific IgE, *A. fumigatus* cutaneous reactivity, bronchial asthma, airway eosinophilia, and bronchiectasis are the major criteria utilized to diagnose APBA. Our data show increased airway eosinophilia, serum IgE, and albumin in the BAL of mice up to 21 days after challenge with *A. fumigatus*, indicating a pathogenic role for *A. fumigatus* long-term persistence. In other published mouse models of ABPA, repeated inoculation is required for these phenotypes to be maintained (45, 50). Further analysis of airway mechanics, host immunology and changes in morphology will be needed to determine the extent of ABPA-disease like pathogenesis in animals with a longer term W72310 infection.

Importantly, ABPA is primarily diagnosed in patients with Cystic Fibrosis, due to an environment in the lung prone to long-term microbial colonization. Mouse models of CF have also shown a strong role for *A. fumigatus* in CF disease pathogenesis. CFTR KO and dF508 mice develop robust inflammatory, IL-4, and IgE (specifically in CD4+ T cells) response to *A.f.* hyphal extract (51). Additionally, exposure of mice to *A. fumigatus* conidia for 24h-72h causes increased inflammation, mucous production and BAL inflammatory cytokine production in CFTR KO compared to WT mice (52) and increased inflammasome activity was observed in CFTR KO mice in response to *A. fumigatus* conidia exposure (53). Future experiments exposing genetically engineered CF mice to W72310 could provide valuable insight into mechanisms of disease pathogenesis and therapeutic efficacy. These types of studies could be especially critical because the concept that strains of *A. fumigatus* can survive in the lung without germinating but cause ABPA-like disease would significantly affect the types of treatments a patient would receive, including antifungals. Moreover, identification of specific fungal alleles that promote long term persistence may help diagnose chronic infections in lieu of transient colonization that would help guide antifungal deployment in these at-risk groups, such as CF patients.

## ACKNOWLEDGEMENTS

This work was supported by the efforts of R.A.C through funding by NIH National Institute of Allergy and Infectious Diseases (NIAID) (grant nos. R01AI130128 and R01AI146121), a pilot award from the Cystic Fibrosis Foundation (CFF) Research Development Award (STANTO15RO), and a CFF research award (CRAMERGO19). C.H.K. was supported by the NIH NIAID Ruth L. Kirschstein National Research Service Award (no. F31AI138354). Core facility support provided by NIH grant P30-DK117469 and NIH grant P20-GM113132 to the Dartmouth BioMT COBRE. J.E.S. is CIFAR fellow in the program Fungal Kingdom: Threats and Opportunities. Computational analyses were performed on the University of California-Riverside HPCC supported by grants from the National Science Foundation (MRI-1429826) and NIH (S10OD016290). T. M. H. received support from NIH grants R01 AI093808 and R01 139632 and Core Grant P30 CA008748 (to MSKCC). J. J. O. received support from NIH grant R01 Al139133.

---

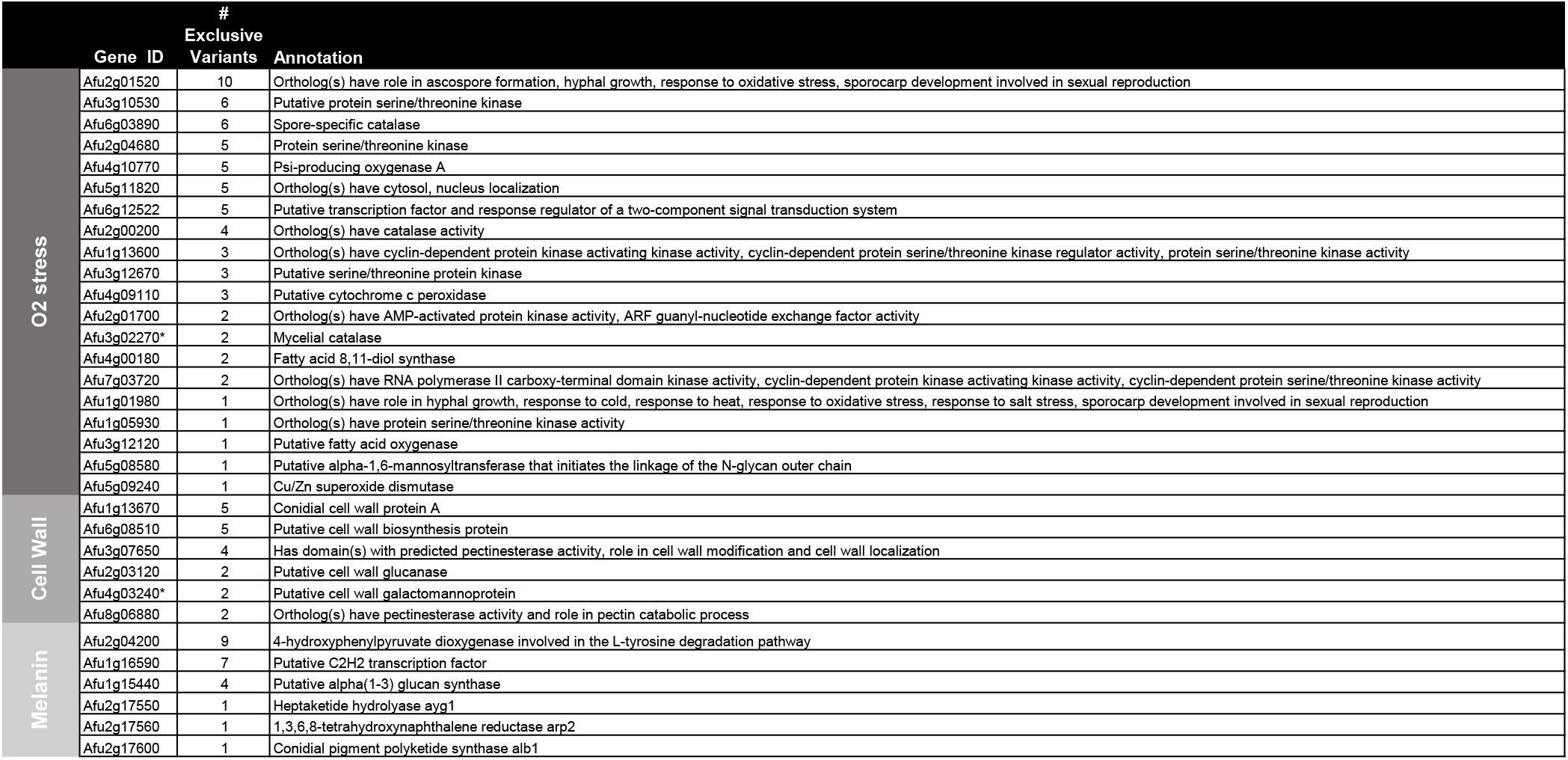
Non-synonamous variants found in W7 relative to AF293, that are not found in CEA10.

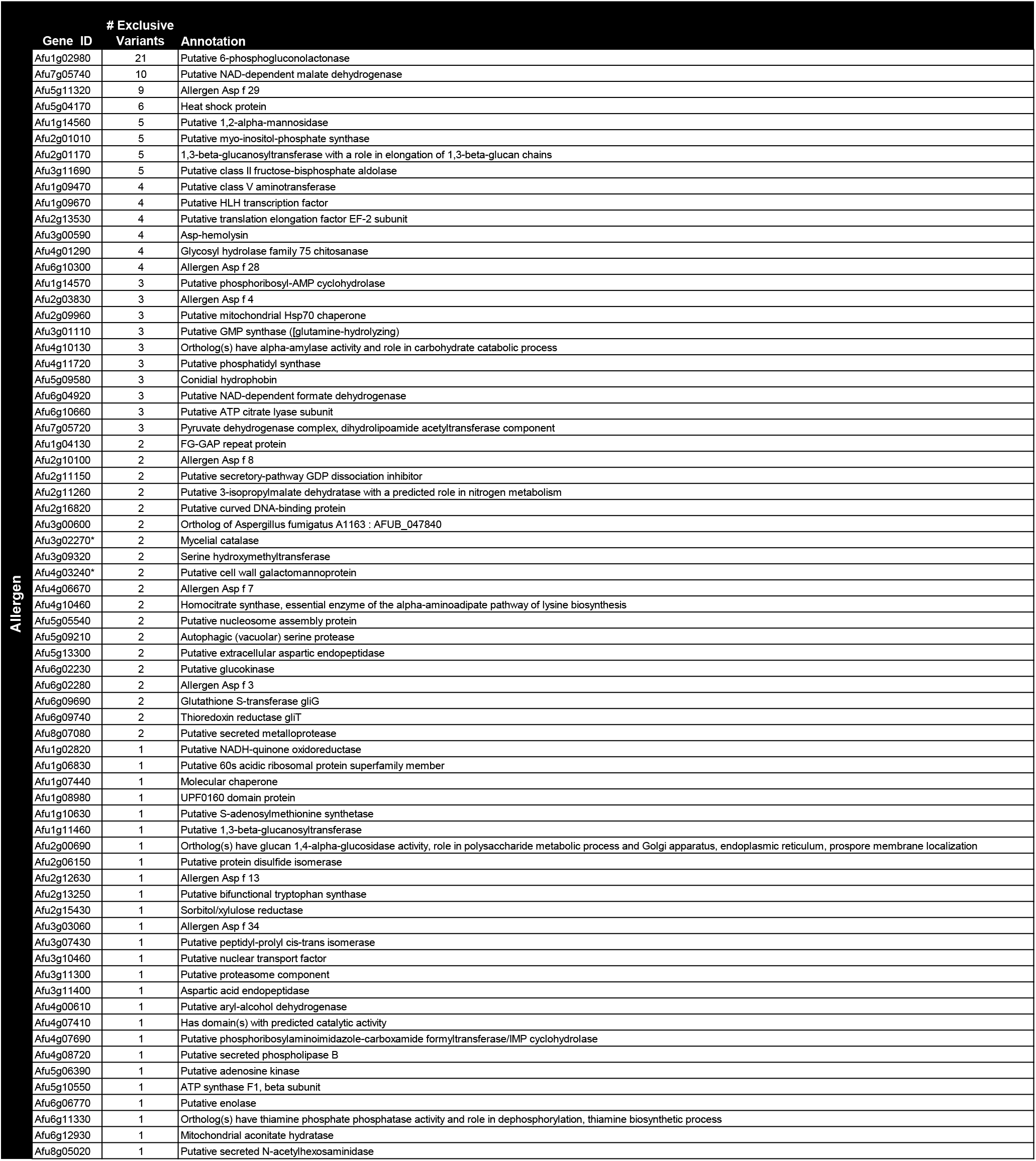
Non-synonamous variants found in W7 relative to AF293, that are not found in CEA10.

**Supplemental Figure 1:**
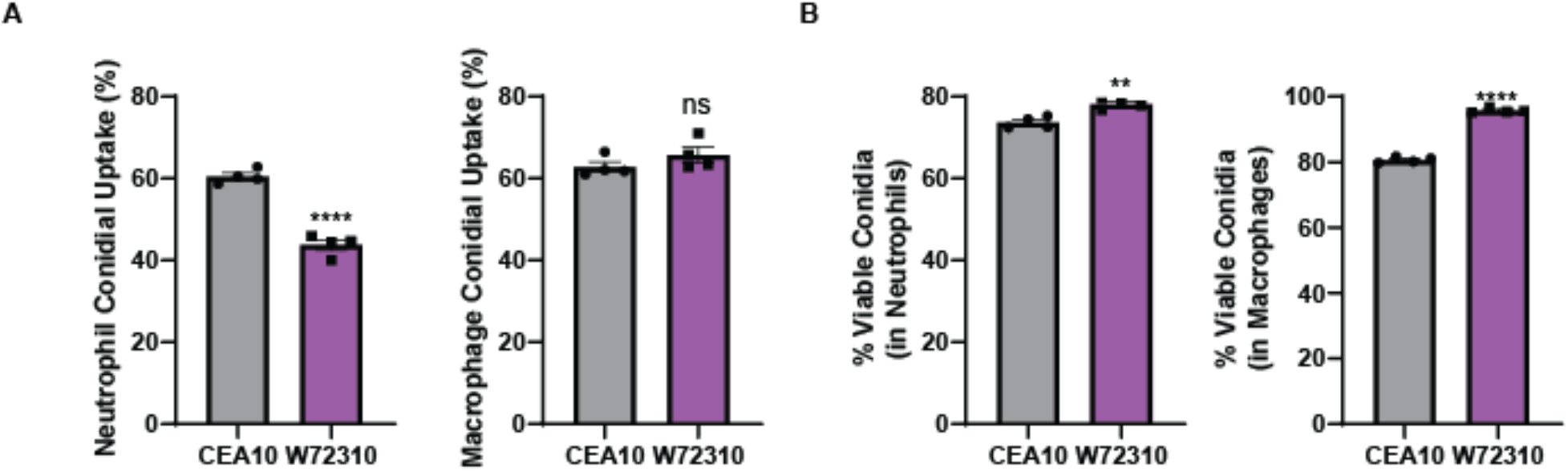
W72310 is more resistant to bone marrow cell-induced death than CEA10. **(A)** AF633-stained mRFP-CEA10 and mRFP-W72310 live conidia were incubated with murine bone marrow for 16 hours and analyzed by flow cytometry for % positive conidia (AF633) in neutrophils and macrophages and **(B)** % viable conidia (AF633^+^/RFP^+^) in neutrophils and macrophages. Data include 3 independent experiments with 3-4 replicates per experiment. Mann-Whitney with Dunn’s multiple comparison were used and NS>0.05; * P≤0.05; ** P≤0.01; *** P≤0.001; **** P≤0.0001.

